# Co-infection with *Streptococcus* and *Rothia* spp. drives prophage dynamics in *Pseudomonas aeruginosa* in an artificial sputum model

**DOI:** 10.1101/2025.05.13.653867

**Authors:** Laura L Wright, Adrian Cazares, Wendy Figueroa, Laura Chatterley, Thomas Willmott, Siobhan O’Brien, Joanne L Fothergill

## Abstract

**Objectives:** Chronic bacterial lung infections with *Pseudomonas aeruginosa* are common in people with cystic fibrosis (CF) but interactions with other commonly seen members of the CF lung community, such as *Streptococcus* and *Rothia* spp., are not well understood. The aim of this study was to determine the impact of other species on *P. aeruginosa* using an artificial sputum medium (ASM) model designed to mimic the conditions in the CF lung.

**Methods:** *P. aeruginosa* LESB58, representing a CF epidemic strain, was inoculated into ASM either alone or in co-culture with *Streptococcus anginosus, Streptococcus vestibularis, Rothia mucilaginosa* or *Rothia dentocariosa*. Gene expression was assessed using RNAseq at 72 and 96 h timepoints. Free phage numbers were assessed by plaque assay and qPCR.

**Results:** Differential expression of *P. aeruginosa* genes, including those related to virulence factors such as prophage, type III secretion, exopolysaccharides and pyochelin production, was seen in co-culture conditions compared with single-species culture. In particular, at 72 h, there was lower expression of most LES prophage genes under co-culture conditions, and this corresponded with lower overall free phage at the same timepoint. Specifically, during co culture, there was a reduction in the levels of a D3112-like transposable phage, previously shown to have the ability to drive CF-like adaptations in an experimental evolution model, and an increase in levels of Pf1-like filamentous phage, associated with altered biofilm formation and immune responses.

**Conclusion:** Gene expression changes during co-culture with *Streptococcus* and *Rothia* spp. were identified affecting key *P. aeruginosa* virulence factors including phage. Further work indicated changes to free phage levels of two important *P. aeruginosa* LESB58 prophage with the potential to affect *P. aeruginosa* adaptation, antibiotic resistance and host immune response in the CF lung. This study emphasises the importance of understanding the influence of lung commensals on focal pathogens in a complex microbiome.

## Introduction

People with cystic fibrosis (CF) have defective ion transport across epithelial surfaces in the lung, resulting in a thick sticky mucus forming and reduced clearance of bacteria. Chronic bacterial lung infections are therefore common in people with CF, with *Pseudomonas aeruginosa* being the predominant pathogen in adults and a leading cause of mortality [1]. Recent metagenomics and extended culture studies have revealed that complex microbial communities exist in the lungs of people with CF. These communities include commonly recognised pathogens such as *P. aeruginosa* cohabiting with a variety of other bacterial species [2-7]. Some of the most prevalent genera detected in the CF lung include *Streptococcus* and *Rothia* spp., but little is known about how these organisms may influence the behaviour of known pathogens such as *P. aeruginosa*.

The presence of a high proportion of *Streptococcus* species has been associated with clinical stability of CF lung disease [8]. The most numerically abundant *Streptococcus* species isolated from the lungs of people with CF include those belonging to the *Streptococcus anginosus* (also including *S. intermedius* and *S. constellatus*) and *Streptococcus salivarius* groups (including *S. vestibularis* and *S. thermophilus*) [8-11]. Several studies have reported a correlation between the presence of *S. anginosus* group species and the onset of pulmonary exacerbations [12-14]. Sibley et al. reported co-isolation of *P. aeruginosa* from 73% of *S. anginosus* group positive sputum samples [11], suggesting that important interactions could occur between the species. It has been shown in laboratory studies that the presence of *S. anginosus* can drive changes to *P. aeruginosa* virulence. For example, increased pyocyanin and elastase production in static biofilms, and decreased survival in a *Galleria* model, have been reported when the species are cultured together [15, 16].

Both *Rothia mucilaginosa* and *Rothia dentocariosa* have also been frequently detected in CF sputum [2-4]. A recent study found *R. mucilaginosa* in 83% of a CF cohort, most often alongside *P. aeruginosa* [17], and the presence of *R. mucilaginosa* has been associated with declining lung function [18]. Laboratory studies have suggested that *R. mucilaginosa* may provide *P. aeruginosa* with precursors for metabolites such as glutamate [19].

The transmissible Liverpool Epidemic Strain (LES) is the most common *P. aeruginosa* lineage isolated from people with CF in the UK and has also been reported in Canada [20]. The LES has been associated with greater morbidity [21, 22], and its genome harbours genomic islands and prophages implicated in its ability to establish respiratory infections [23, 24].

In this study, we used an artificial sputum medium (ASM) model, designed to mimic the conditions found in the CF lung, to investigate how *Rothia* and *Streptococcus* species influence the behaviour of the *P. aeruginosa* LES isolate LESB58. We report the effects of co-culture on growth and, using RNAseq analysis, *P. aeruginosa* gene expression. We identified downregulation of genes in prophages associated with the establishment of infection and adaptation of *P. aeruginosa*, but upregulation of genes of a filamentous phage linked to altered host immunology. These changes could also be seen in free phage levels and demonstrate the impact that growth in a polymicrobial environment can have on *P. aeruginosa* biology.

## Materials and methods

### Bacterial strains and growth conditions

ASM was prepared as described by Kirchner et al. [25] and was sterilised by autoclaving. *P. aeruginosa* LESB58, a representative of a CF transmissible lineage [23], was inoculated into 2 mL ASM in 24 well plates either alone or in co-culture with *S. anginosus, S. vestibularis, R. mucilaginosa* or *R. dentocariosa. Streptococcus* and *Rothia* clinical strains were obtained from the nasal tract of healthy individuals [26]. Initial total bacterial inoculum was kept constant at 1 x 10^5^ bacteria per 2 mL culture. For dual species culture 5 x 10^4^ bacteria of each species were inoculated. Cultures were incubated at 37°C with slow shaking (150 rpm) for up to 168 h. All experiments were carried out in triplicate.

### Bacterial growth assays

Cultures at each timepoint were homogenised with an equal amount of sputasol (Oxoid, Basingstoke, UK). Dilution series were plated on *Pseudomonas* selective agar (Oxoid) and grown aerobically at 37°C for 24 h to obtain *P. aeruginosa* counts, or plated onto Colombia agar supplemented with 5% (v/v) horse blood (TCS biosciences, Buckingham, UK) and *Streptococcus* selective supplement (Oxoid) and grown anaerobically at 37°C for up to 72 h to obtain *Streptococcus* and *Rothia* spp. counts. Means were compared for each time point by one-way ANOVA with a Tukey HSD post hoc analysis using SPSS software (v 25.0).

### RNA Extraction and RNAseq Library Preparation

Cultures were removed from incubation at the appropriate timepoint and 6 mL trizol was immediately added to the 2 mL culture and mixed well. RNA was then extracted using the Zymo Directzol RNA miniprep kit (Zymo, Irvine, USA) following the manufacturer instructions. RNA samples were treated with the Turbo DNA-free kit (Invitrogen, Waltham, USA), and the absence of any remaining *P. aeruginosa* DNA was confirmed by a negative PCR. RNA concentration and purity were determined by using Qubit and nanodrop, respectively.

RNAseq library preparation and sequencing were carried out by the Centre for Genomic Research at the University of Liverpool. RNA was depleted with the RiboZero Magnetic Kit for bacteria (Illumina, San Diego, USA) and depletion confirmed with a Bioanalyzer RNA pico-chip. RNA-Seq libraries were then prepared using the NEBNext Ultra Directional RNA Library Prep Kit for Illumina, with 14 cycles of amplification, and purified using AMPure XP beads. Libraries were quantified using Qubit and size distribution assessed with the Agilent 2100 Bioanalyser. Final libraries were pooled in equimolar amounts and the pools assessed for quantity and quality using the Bioanalyser and by qPCR using the Kapa Illumina Library Quantification Kit (Roche, Basel, Switzerland) on a Roche Light Cycler LC480II. RNA libraries were sequenced on the Illumina HiSeq 4000 platform with version 1 chemistry to generate 2 x 75 bp paired-end reads.

### RNAseq Data Analysis

Basecalling and de-multiplexing of indexed reads was performed by CASAVA version 1.8.2 (Illumina). The resulting raw FASTQ files were trimmed to remove Illumina adapter sequences using Cutadapt version 1.2.1 [27]. The option “-O 3” was set, so that the 3’ end of any reads which matched the adapter sequence over at least 3 bp was trimmed off. The reads were further trimmed to remove low quality bases, using Sickle version 1.200 with a minimum window quality score of 20. After trimming, reads shorter than 20 bp were removed (over 99.8% of reads survived this step across all samples).

Reads were aligned to the LESB58 genome using Tophat version 2.1.0 [28]. The genome reference, along with gene annotations, used for *P. aeruginosa* LESB58 mappings was downloaded from the Pseudomonas Genome DB database (https://www.pseudomonas.com). The option for read mat orientation was set as “--library-type fr-firststrand”, and the option for number of hits to be reported was specified as “-g 1”, which instructs the mapper report at most one alignment which shows the best mapping quality. The resulting read mapping percentages were high, ranging from 91.37% to 96.88%.

Gene expressions were calculated from the alignment files using htseq-count [29]. The raw count data were converted into FPKM (Fragments per Kilobase per Million reads) values for initial visualisation and overall gene expression exploration. The raw count numbers per gene were also used during the subsequent differential expression analysis. The main processes of the analysis included data variation assessment, data modelling, model fitting, testing and DE (DifferentiallyExpressed) gene detection. Briefly, variation between samples and sample groups was assessed via principal component analysis (PCA); count data variation was modelled using a generalized linear model (GLM), with the mean expression of each sample group representing a different parameter in the model; after parametrisation, GLM was used to obtain the log2 Fold Change (log2FC) values for the required comparisons; common, trended and tag-wise dispersion parameters were estimated and the latter was used for significance testing; the estimated log2FC were tested using a Likelihood-Ratios (LR) test and P-values associated with log2FC were adjusted for multiple testing using the False Discovery Rate (FDR) approach; significantly differentially expressed genes were defined as those with FDR-adjusted P-value < 5%. All the DGE (Differential Gene Expression) analyses were performed in R (version 3.3.3) environment using the DESeq2 [30] package.

### Plaque assays

Cultures in ASM were grown at 37°C with slow shaking for 24 and 72 h. Following the incubation period, the cultures were homogenised with 1 mL of sputasol (Oxoid), and filter-sterilised (0.22 µm Millipore membrane). Serial dilutions were plated onto *Pseudomonas* selective agar to determine LESB58 colony forming units (CFU/mL). In order to quantify the free phage produced in each condition, bacterial lawns of the strain PA□ (a *P. aeruginosa* strain highly susceptible to phage infection [31]) were made by mixing 150 µL of an overnight culture and 3.5 mL of top agar (0.7% LB agar) and plating them on Luria-Bertani (LB) plates. Serial dilutions of the polymicrobial cultures were then spotted onto the lawns to determine the plaque forming units per mL (PFU/mL). The production of free phage was calculated as the number of plaques produced per colony forming unit (PFU/CFU) and then normalised to the values of LESB58 alone in each time point. The experiment was performed with 6 replicates. Welch’ T-tests were performed followed by a Bonferroni correction using R package EnvStat (R-4.0) to detect statistically significant differences between the different polymicrobial conditions and the control (LESB58 alone).

### DNA extraction and phage qPCR

DNA was extracted from the triplicate ASM cultures stored in trizol reagent (Invitrogen) using a phenol/chloroform method according to the manufacturer’s protocol [32]. The resulting DNA pellet was resuspended in 100 μL molecular grade water at 65 °C. DNA was diluted 1:10 before use in qPCR assays. As described previously by James et al [33] primers targeting total phage (free phage + prophage) and prophage alone were used to determine free phage abundance for LESB58 phage 2-6 by qPCR and bacterial density was calculated using primers specific to *P. aeruginosa*. Standards for each primer set were prepared by conventional PCR and a dilution series of the appropriate standard was included in each qPCR run on the Rotorgene cycler (Qiagen, Hilden, Germany). Data were analysed using the Rotorgene Q software to determine copy numbers and phage to bacterium ratios were calculated. Means were compared by independent samples T-test using SPSS Statistics (v25.0) software.

## Results

### Growth rates in ASM

No significant differences in *P. aeruginosa* growth rate were seen in any co-culture condition compared with single culture (P>0.05) (Figure 1). However, both *Streptococcus* species, along with *R. mucilaginosa*, showed reduced growth when co-cultured with LESB58. *S. vestibularis, S. anginosus* and *R. mucilaginosa* survived to 168 h in ASM when cultured alone, but growth was significantly reduced from 72 h onwards when cultured together with *P. aeruginosa* (P<0.001). In contrast, *R. dentocariosa* growth was significantly increased at 48 h under co-culture conditions compared with single culture (P<0.001), although the species was not detectable by the 72 h timepoint in either single or co-culture. The limit of the co-culture before *P. aeruginosa* outcompetes is 72 hours.

**Figure 1.**
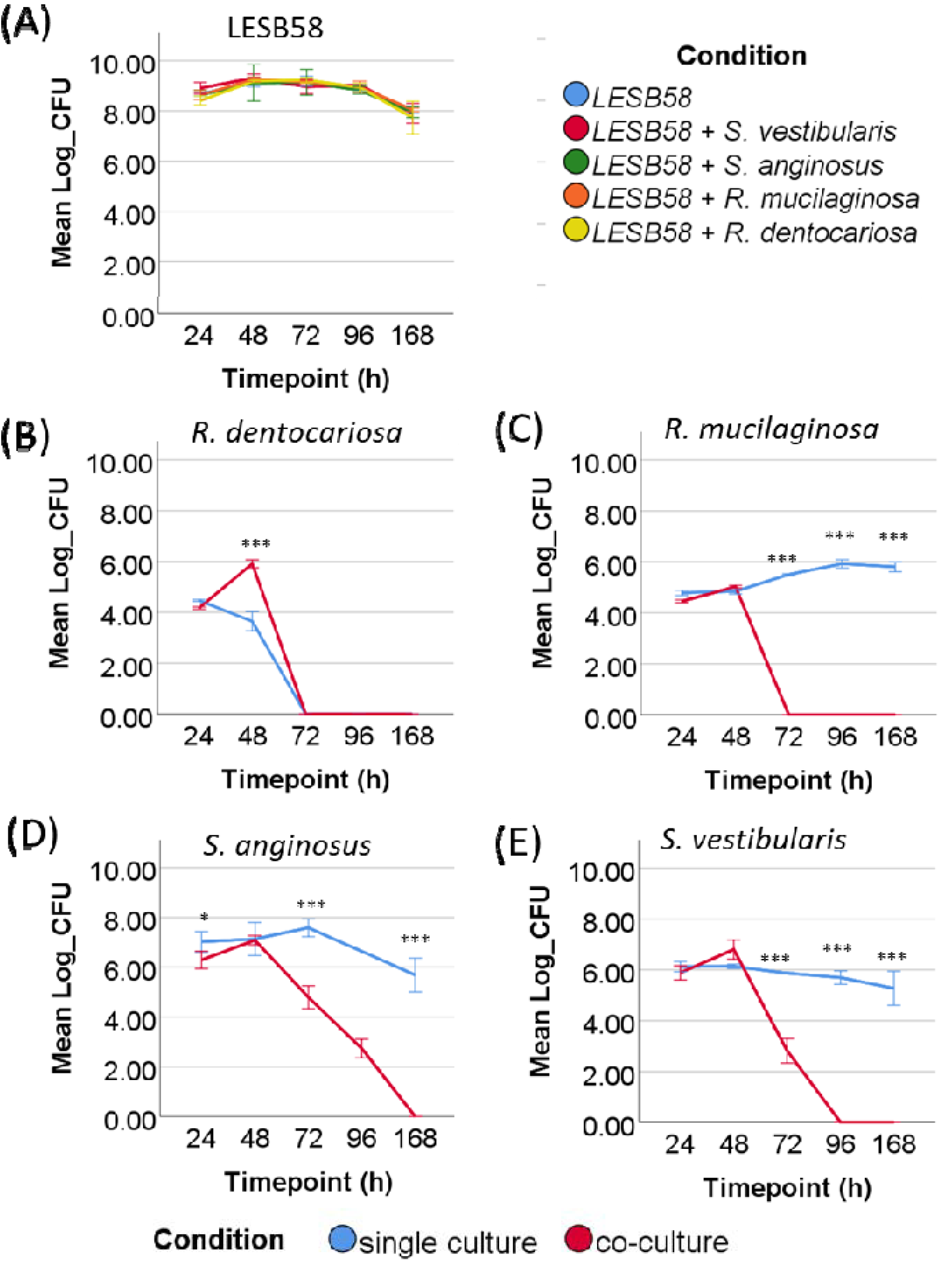
Growth rate of *P. aeruginosa, Streptococcus* and *Rothia* species in single and co-culture conditions when grown in artificial sputum medium. (A) *P. aeruginosa* LESB58 growth rate in single and co-culture conditions. (B) *R. dentocariosa* (C) *R. mucilaginosa* (D) *S. anginosus* (E) *S. vestibularis* growth rates in single (blue) and co-culture with *P. aeruginosa* LESB58 (red). Error bars indicate ±1 SD, ^*^ P<0.05,^**^ P<0.01, ^***^ P<0.001.

### *P. aeruginosa* gene expression changes in co-culture

To assess gene expression changes in co-culture, RNA was extracted from ASM cultures of *P. aeruginosa* alone and in co-culture at 72 h and 96 h timepoints representing mid and late stationary phases of growth respectively. In total four different co-culture conditions were investigated by growing *P. aeruginosa* LESB58 with *S. anginosus, S. vestibularis, R. mucilaginosa* or *R. dentocariosa*. The resulting gene expression data can be seen in Supplementary Table 1.

Initial analysis using principal components analysis (Figure 2) indicated distinct *P. aeruginosa* gene expression patterns across conditions at 72 h. Co-culture with *S. anginosus* and *S. vestibularis* clustered closely, whereas co-culture with *R. mucilaginosa* clustered the furthest from all other conditions. Overall, co-culture with *S. vestibularis* and *R. mucilaginosa* displayed the most divergent expression patterns compared to LESB58 growing alone. At the 96 h time point, the different groups clustered more closely together, with the exception that the co-culture with *S. vestibularis* clustered separately and closer to the 72 h culture conditions. Since the majority of the differences were seen at 72 h, and growth rate data indicated that populations of the *Streptococcus* and *Rothia* species were declining after 72 h, this time point became the focus of further analysis to determine which genes were responsible for the different gene expression patterns.

**Figure 2.**
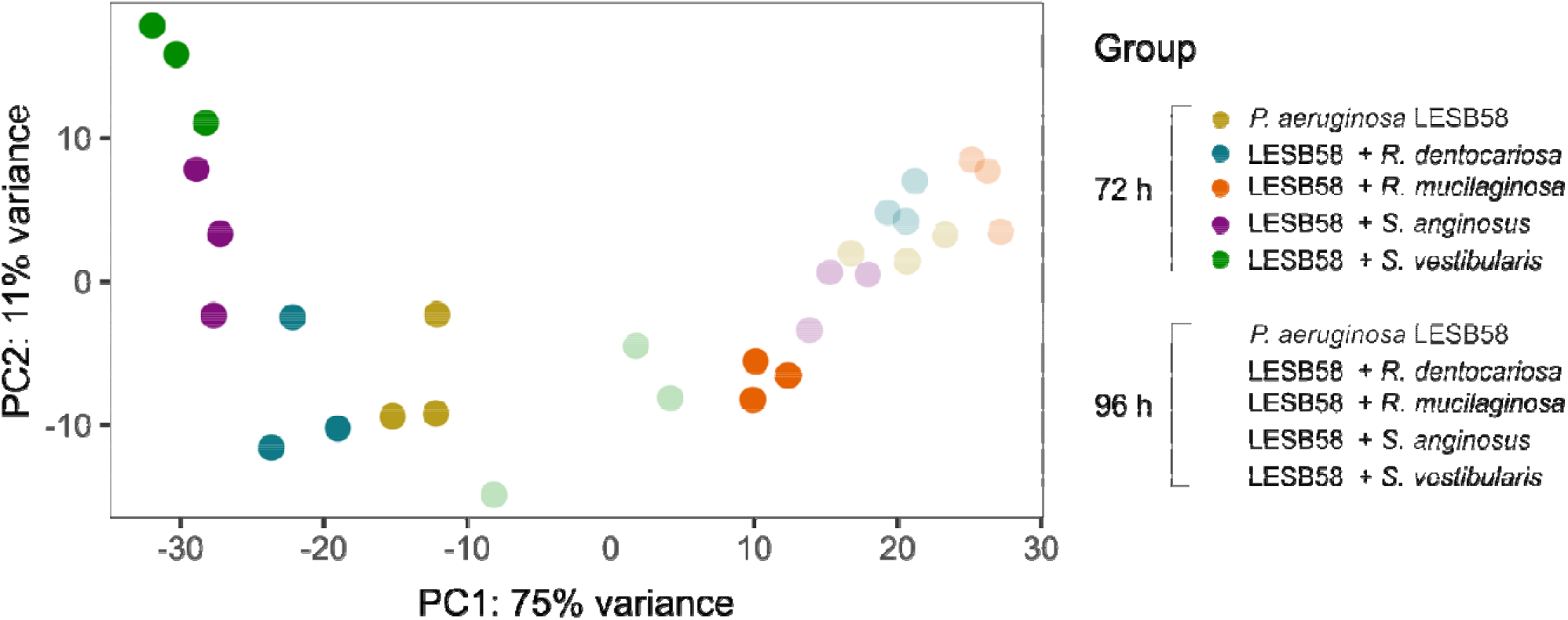
Principal component analysis of gene expression data from RNAseq of *P. aeruginosa* grown either alone or co-cultured with either *Rothia dentocariosa, Rothia mucilaginosa, Streptococcus anginosus* and *Streptococcus vestibularis*.

Genes were initially filtered on a false discovery rate (FDR) of < 0.05 and this indicated that a large proportion of the 5927 genes in the *P. aeruginosa* LESB58 genome was significantly differentially expressed at 72 h in each co-culture condition (*R. dentocariosa* 1185, *R. mucilaginosa* 3176, *S. anginosus* 2329, *S. vestibularis* 2983. Therefore, in order to consider only the most differentially expressed genes, a further filter of log2 fold change (FC) values of < -1.5 or > 1.5 was applied to the data (Figure 3).

**Figure 3.**
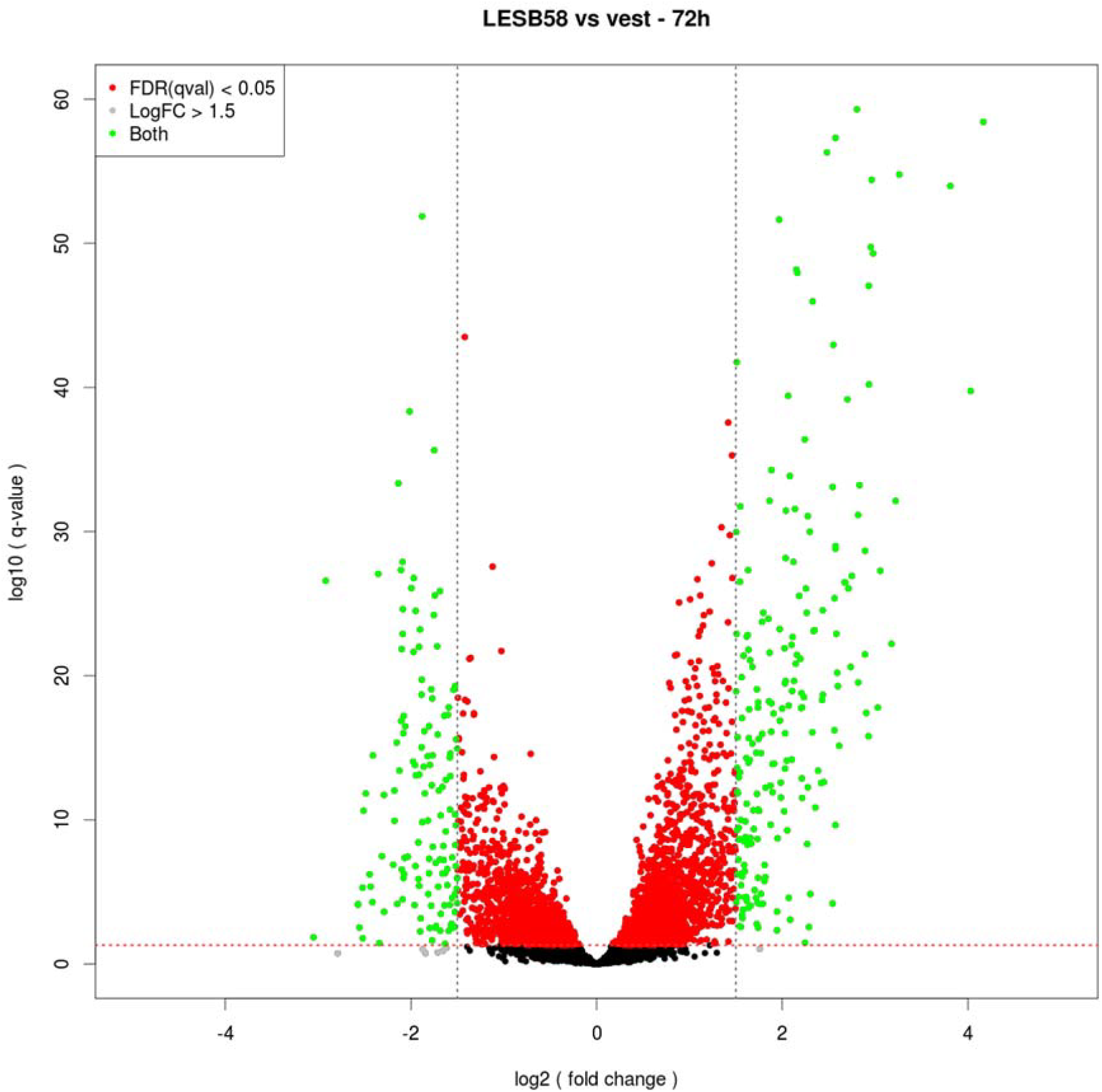
Distribution of False Discovery Rate (FDR) vs Log2 Fold Change (FC) values for LESB58 co-cultured with *S. vestibularis*. Cut offs for FDR < 0.05 (horizontal line) and Log2FC ±> 1.5 (vertical lines) are indicated. Only genes that reached both of these cutoffs, indicated in green, were taken forward for further analysis.

After these criteria were applied (FDR <0.05 and log2FC < -1.5 or > 1.5), a total of 553 genes that were differentially expressed in one or more of the co-culture conditions were identified. The highest number of differentially expressed genes was seen when *P. aeruginosa* was co-cultured with *S. vestibularis* (351 genes), followed by co-culture with *R. mucilaginosa* (200), *S. anginosus* (140) and *R. dentocariosa* (19) (Figure 4).

**Figure 4.**
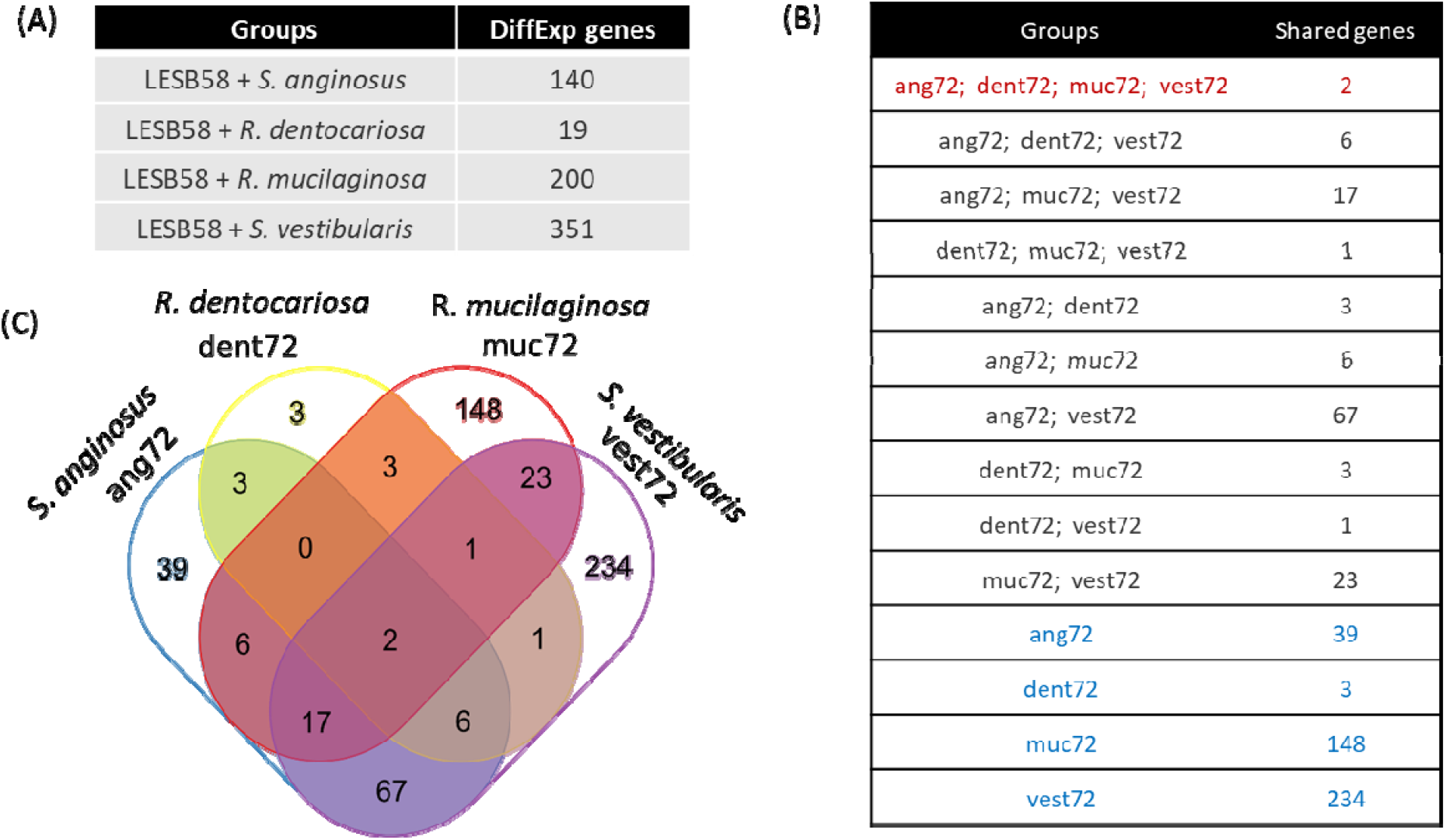
(A) Table showing the numbers of differentially expressed *P. aeruginosa* LESB58 genes at 72 hours in co-culture with either *S. anginosus, S. vestibularis, R. mucilaginosa* or *R. dentocariosa* in ASM when compared to single species culture. Venn diagram (B) and table (C) indicate the numbers of shared genes between different co-culture conditions. Genes were filtered based on a false discovery rate (FDR) of < 0.05 and log2FC values of < -1.5 | > 1.5.

At 72 hours (Day 3) a total of 234, 148, 39 and 3 unique *P. aeruginosa* LESB58 genes were differentially expressed in co-culture with *S. vestibularis, R. mucilaginosa, S. anginosus*and *R. dentocariosa*, respectively. A further 67 and 3 genes were only differentially expressed in both members of the *Streptococcus* and *Rothia* spp., respectively. The largest number of unique differentially expressed genes were seen in co-culture with *S. vestibularis* with 234 genes where 120 were at higher levels of expression, while 114 were lower compared with *P. aeruginosa* single culture.

### Functions linked to differentially expressed genes

The genes that were differentially expressed are involved with a variety of functions, including metabolism, transcriptional regulation, type III secretion, pyochelin biosynthesis, exopolysaccharides and production of LPS. Of those genes that were differentially expressed, 69 had a log2FC value of more than +/-2.5 in co-culture with at least one of the four species and 17 had a log2FC of more than 3. The largest log2FC seen was with PLES_36261, a gene associated with type III secretion, for which expression was log2FC -6.09 lower in co-culture with *S. vestibularis*.

Several of the 69 most differentially expressed genes were located on genomic islands. Three are part of LES-GI 3 (PLES_26251 and PLES_26311, both encoding hypothetical proteins, and PLES_26261, encoding a heavy metal translocating P-type ATPase) and other three are located on LES-GI 5 (PLES_44881, PLES_45031 and PLES_45071, all encoding hypothetical proteins). Some of the genes were also located on LES prophage 1 (PLES_06091, ptrB, a type III secretion repressor), LES prophage 5 (PLES_25611 and PLES_25341, both encoding hypothetical proteins) and LES prophage 6 (PLES_41231, encoding a hypothetical protein).

Expression of the psl exopolysaccharide (*pslO*) gene was higher at the 72 hour timepoint when *P. aeruginosa* was co-cultured with *S. vestibularis* (log2FC of 2.57), *S. anginosus* (2.47) and to a less extent *R. dentocariosa* (1.72), while expression of pslK, was also higher at the 72 hour timepoint for *P. aeruginosa* co-cultured with *S. vestibularis* (1.63) and *R. mucilaginosa* (2.73). The psl locus is involved in the production of Psl polysaccharide, which is an important component of the biofilm matrix with functions including prevention of complement opsonisation, attachment to epithelial cells and initiation of a proinflammatory response.

Expression of the pyochelin (*pchC*) gene, involved in pyochelin biosynthesis, was lower for *P. aeruginosa* cultured with *S. vestibularis* (−2.57). Pyochelin and pyoverdine, are siderophores that can chelate iron and have been shown to be important in the virulence of *P. aeruginosa*.

Several genes associated with type III secretion system (T3SS) were differentially expressed under co-culture conditions. Expression of exsB was higher at 72 hours when *P. aeruginosa* was co-cultured with *S. vestibularis* (3.06) and *S. anginosus* (2.34) compared to single culture. In contrast, expression of the genes PLES_36261 and PLES_36281 was lower under co-culture conditions with *S. vestibularis* (−6.09 and -2.34, respectively) and *R. mucilaginosa* (PLES_36281 only, -2.59). PLES_36261 and PLES_36281 are also known as pcr3 and pcr1; pcr3 is required for type III secretion and pcr1 is a type III secretion system repressor [34]. Expression of ptrB, a repressor of type III secretion located on LES prophage 1, was higher when *P. aeruginosa* was co-cultured with *R. mucilaginosa*.

### Changes in phage gene expression and free phage levels during co-culture

A large proportion of the filtered genes with differential expression in one or more of the co-culture conditions were LES prophage genes (77/553), the vast majority of which were downregulated (73/77). To assess whether more prophage genes were significantly differentially regulated but below the 1.5 log2FC cut off, we assessed gene expression of all genes contained within each of the six prophages that have been identified in LESB58 [23]. We found that a large proportion of genes were significantly down regulated during co-culture conditions at 72 h for phage 2-6, although for phage 1 (a pyocin) and phage 6 (a Pf1-like filamentous phage) there was a mixture of up and down regulated genes. Different expression profiles are shown in Figure 5 (Phage 4 and 6) with similar analysis for the other phage available in supplementary Figures 1-4. Several of the genes that were differentially expressed included *recA, recX* and *lexA*, which are associated with *P. aeruginosa* SOS response [35] (supplementary Table 1), however, the gene expression responses in *P. aeruginosa* were not consistent.

**Figure 5.**
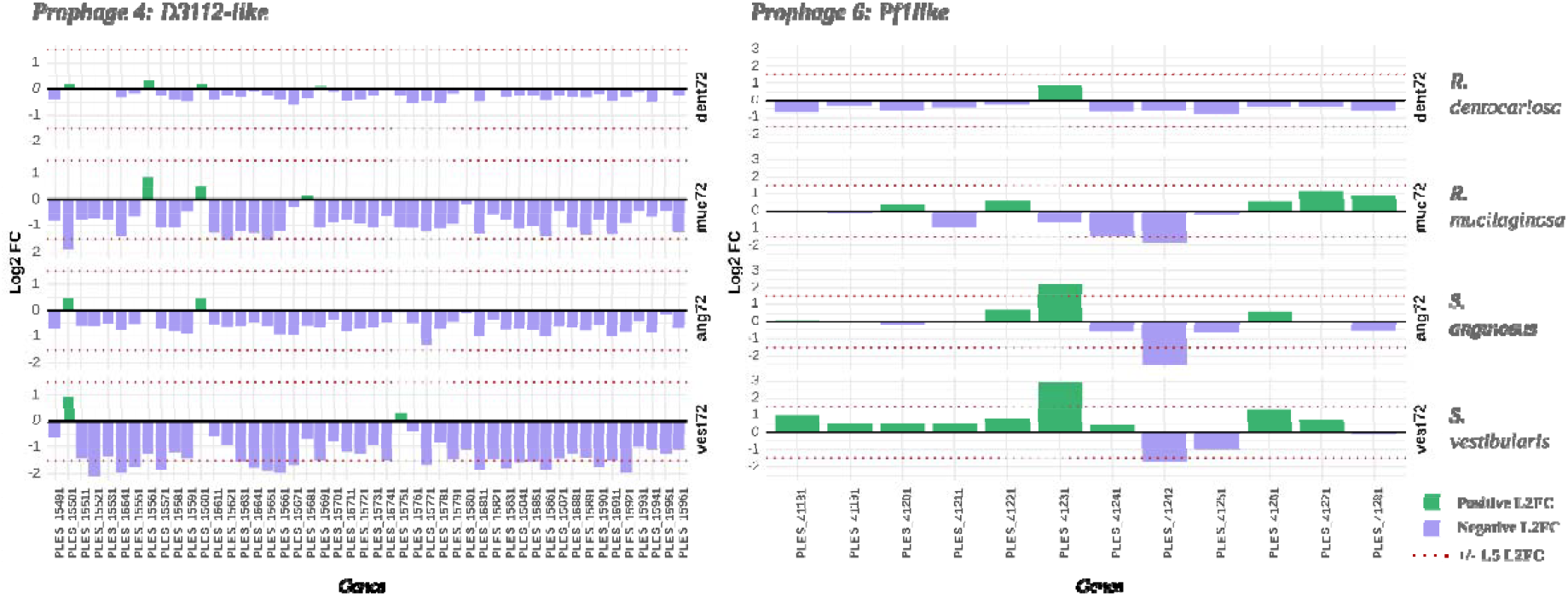
Gene expression changes in *P. aeruginosa* LESB58 D3112-like prophage 4 and Pf1-like prophage 6 at 72 hours in co-culture, compared with single species culture. Upregulated genes compared to *P. aeruginosa* single culture are shown in green and downregulated genes are shown in blue. The ± 1.5 log2FC cut off is indicated with a dotted line.

To examine whether the differential expression of phage genes at 72 h was linked to a reduction in free phage numbers, we assessed prophage induction using plaque assays. Prophage induction was significantly reduced in all of the dual species culture conditions compared to LESB58 culture alone (Figure 6). A similar pattern was seen at a 24 h time point when *P. aeruginosa* was co-cultured with both *R. mucilaginosa* and *S. anginosus*.

**Figure 6.**
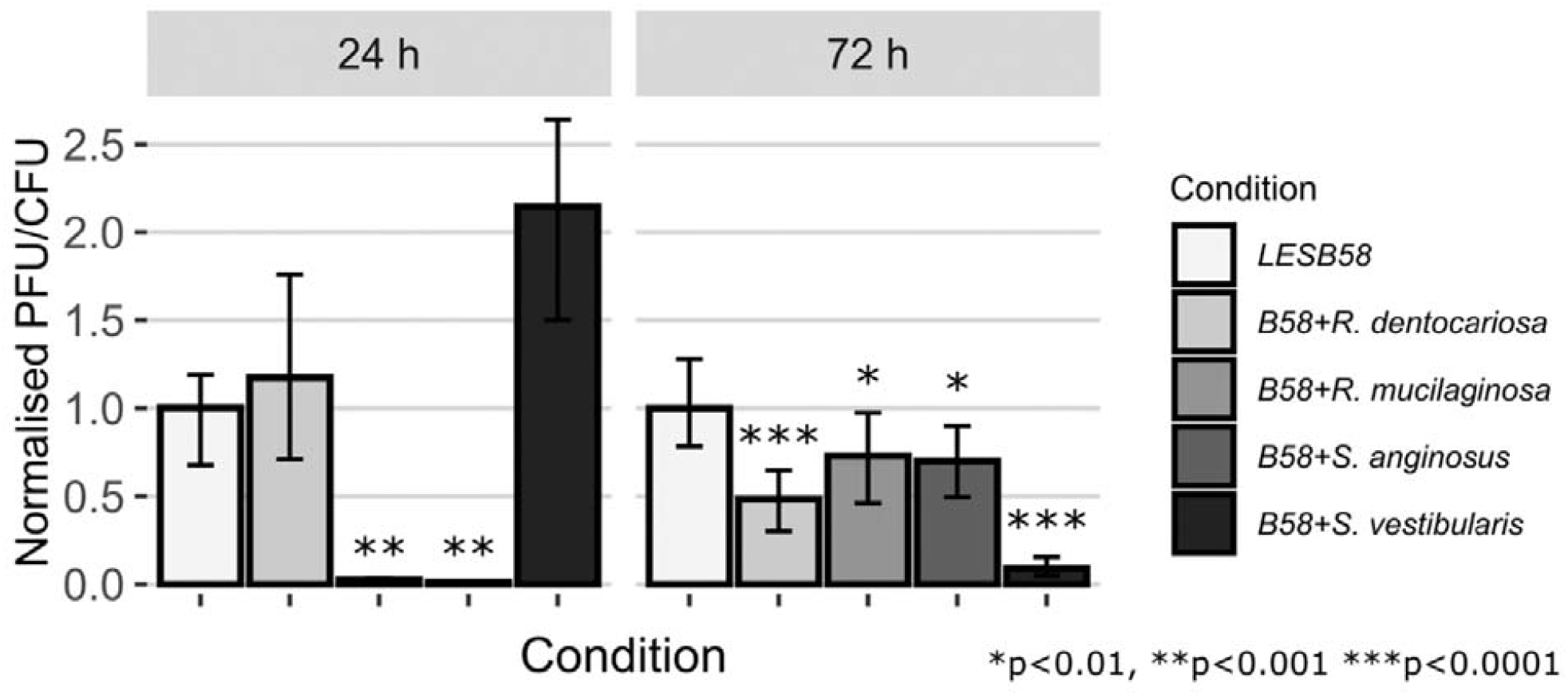
Total free phage levels assessed by plaque assays under different co-culture conditions and timepoints. The production of free phage was calculated as the number of plaques produced per colony forming unit (PFU/CFU) and then normalised to the values of LESB58 alone in each time point. ^*^P<0.01,^**^ P<0.001, ^***^ P<0.0001.

In order to identify the biggest contributors to the reduction in phage levels at 72 h, qPCR was used to assess free phage levels for LESB58 phages 2-6 (Figure 7). Levels of LESB58 phage 4, a D3112-like transposable phage, were lower in all the co-culture conditions compared to LESB58 alone, with this being significantly different when LESB58 was grown with *R. dentocariosa* (P=0.037) and *S. vestibularis* (P=0.031). Levels of phage 6, a Pf1-like filamentous phage, were significantly higher at 72 h for *S. vestibularis* (P=0.050), *R. mucilaginosa* (P=0.019) and *R. dentocariosa* (P=0.015) co-culture conditions compared with LESB58 alone.

**Figure 7.**
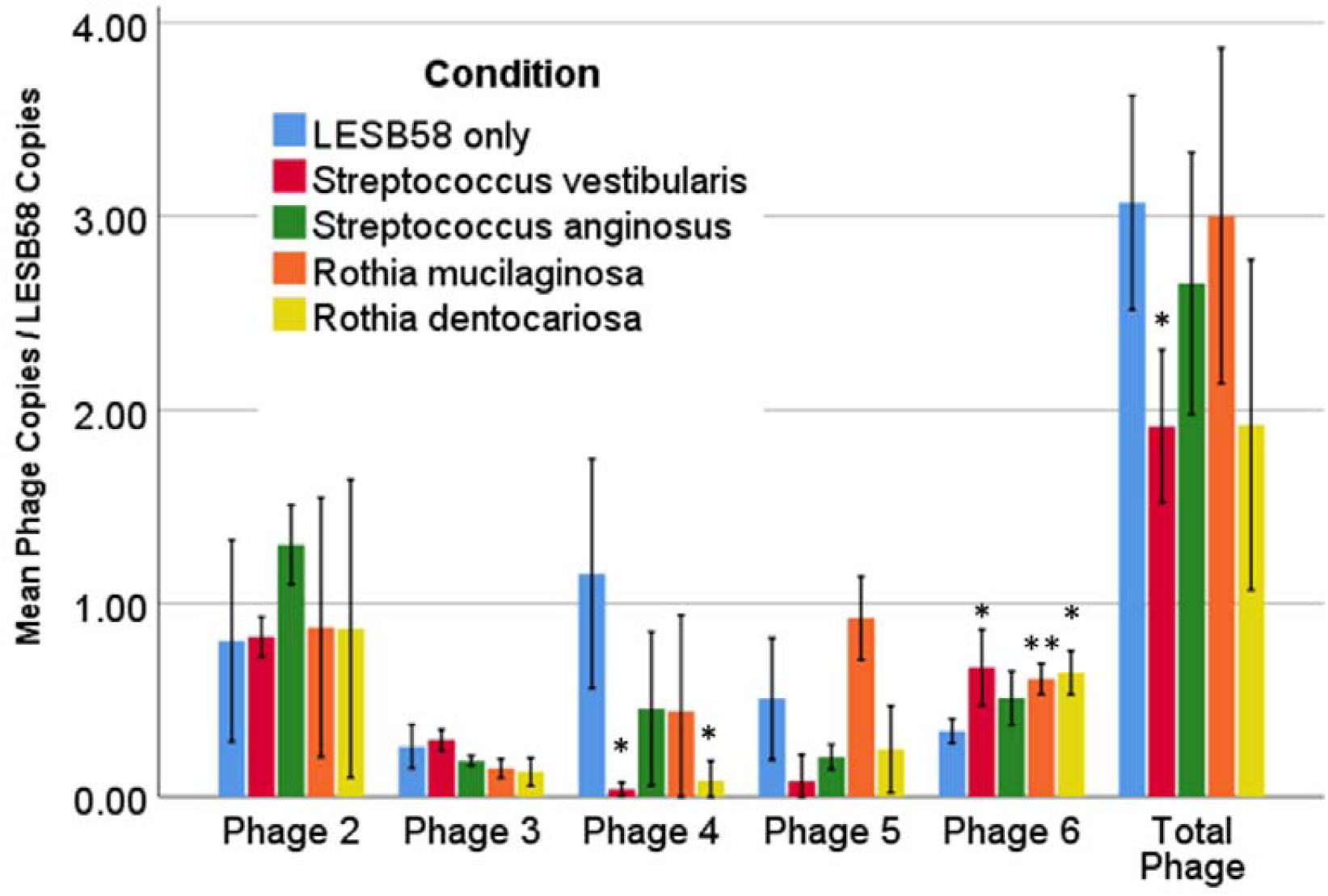
Mean LESB58 free phage levels in single or co-culture conditions. Phage levels determined by qPCR were divided by the calculated LESB58 copy number for each sample to obtain phage to bacterium ratios. Error bars indicate ±1 SD.

## Discussion

Temperate phages are a major influence on *P. aeruginosa* in chronic infection [36] and here we show that their expression is influenced by the wider microbiome. Gene expression of most LESB58 prophage genes was reduced under all dual species co-culture conditions and this corresponded with significant reductions in overall spontaneous phage production.

The majority of *P. aeruginosa* LESB58 phage genes were downregulated under all of the dual species culture condition suggesting a common response across the two genera. Six prophages have been identified in the LESB58 strain and have been shown to affect virulence of *P. aeruginosa* [23]. Specifically, we found that levels of D3112-like transposable LES phage 4 were significantly reduced during co-culture with *R. dentocariosa* and *S. vestibularis* compared with LESB58 single culture. This phage is a transposable phage that has been linked to adaptation and therefore a reduction in the production of this phage may alter the ability of *P. aeruginosa* to adapt rapidly. D3112-like phage have been commonly detected in the sputa of people with CF [37]. They have also been associated with the ability to drive CF-like adaptations, such as mutations in quorum sensing and motility associated genes, in an artificial sputum experimental evolution model [38]. The reduction in the levels of these phage during co-culture in our study suggests that co-culture may slow the development of these adaptations in *P. aeruginosa* biofilms.

On the other hand, levels of the Pf1-like filamentous LES phage 6 were higher under all co-culture conditions compared to LESB58 alone. This phage is a Pf-like phage and the presence of this phage has been linked to alterations in biofilm architecture, antibiotic resistance and host immune responses. Pf phage have been detected in CF sputa [39] and upregulation of several of the Pf1-like prophage genes in PA01 has also been reported in *P. aeruginosa* biofilm models [40, 41]. Pf phage have an important role in both biofilm formation and host immunity [42]. They contribute to cell lysis and the release of extracellular DNA, an important stabilising component of biofilms [42, 43]. In addition, the phage can themselves form a liquid crystal structure together with other components of the biofilm matrix resulting in a more stable biofilm structure [39, 43]. This is associated with increased antibiotic resistance both through reduced penetration of antibiotics through the biofilm and sequestration of cationic antibiotics such as aminoglycosides by the negatively charged phage [39, 44]. Production of Pf phage has been shown to lead to a less invasive and more chronic infection phenotype, characterised by a reduced inflammatory response and suppression of bacterial clearance by phagocytosis, in the context of both chronic CF lung infection and wound infection models [39, 45, 46].

Overall, this study has utilised sequencing technologies in order to characterise interactions within the microbial community present in complex clinical niches such as chronic lung infections. Bacteria previously described as commensals were shown to alter key features of *P. aeruginosa* biology including phage production. Production of a transposable phage was decreased and could impact bacterial adaptation and evolution in the lungs. Filamentous phage production was increased and these phages have been associated with altered biofilm formation, antibiotic resistance, chronic infection and altered immune responses. *P. aeruginosa* infection biology is complex, particularly in the context of the altered environment of the CF lung. A better understanding of interactions between species in the CF lung may uncover a deeper understanding of adaptation, evolution and potentially new therapeutic targets.

## Acknowledgments

We acknowledge the Medical Research Foundation [Grant MRF-091-0006-RG-FOTHE] and the CF Trust and CF Foundation (SRC022) for funding.

## Author contributions

LW – methodology; validation; formal analysis; investigation; data curation; visualization; writing – original draft. AC – methodology; validation; formal analysis; investigation; data curation; visualization; writing – original draft. WF – methodology; validation; formal analysis; investigation; data curation; visualisation; writing – original draft. LC – investigation; writing – reviewing, editing. TW – visualization; data curation; writing – original draft. SB – conceptualization; writing – reviewing, editing; funding acquisition. JF – conceptualization; methodology; writing – reviewing, editing; supervision; project administration; funding acquisition.

## Data availability

The data has been deposited in the NCBI BioProject database (BioProjectID: PRJNA1247576).

