## Supplementary Data for "Co-infection with *Streptococcus* and *Rothia* spp. drives prophage dynamics in *Pseudomonas aeruginosa* in an artificial sputum model"

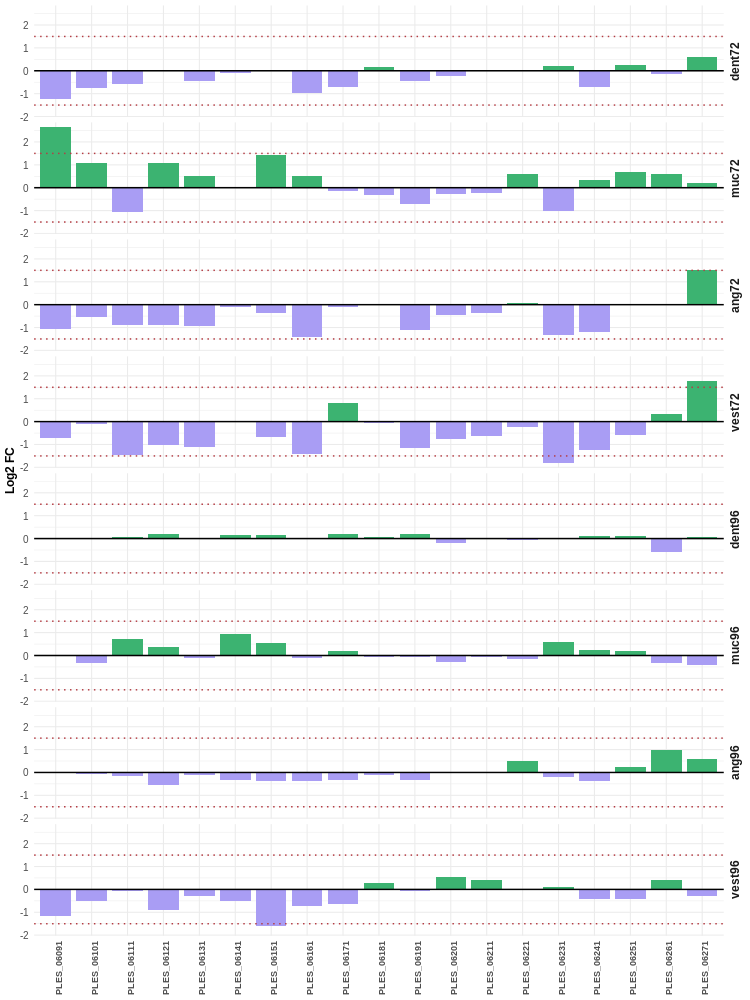

**Supplementary Figure 1.** Prophage 1: Pyocin R2. Log2FC values distribution of LESB58 prophage genes during inter-species bacterial interactions.

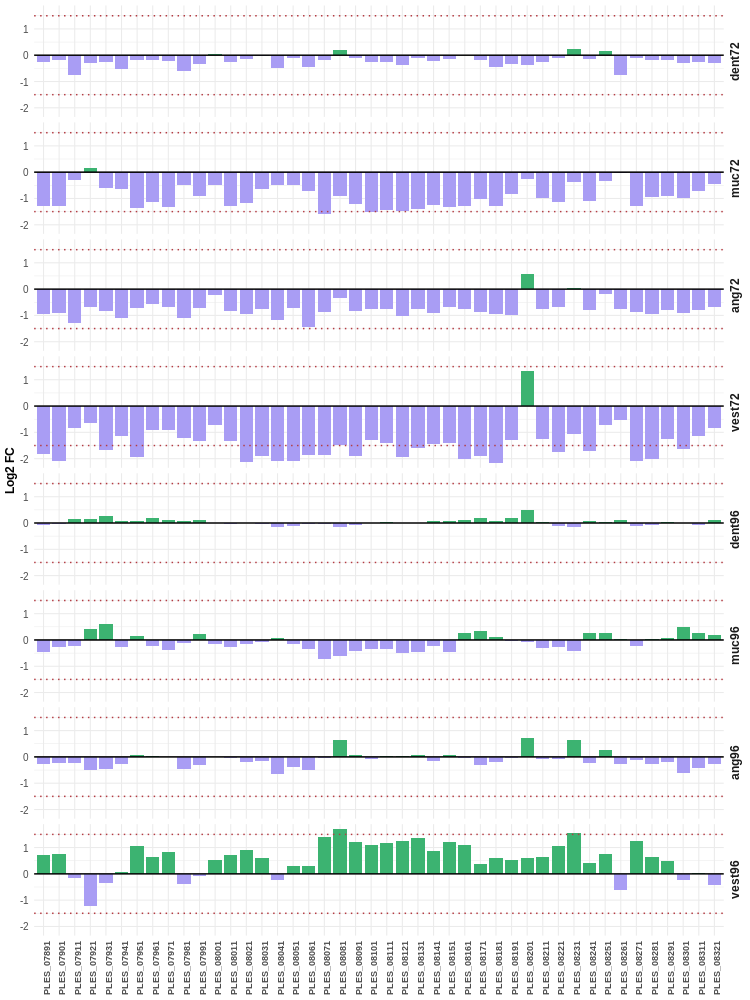

**Supplementary Figure 2.** Prophage 2: F10-like. Log2FC values distribution of LESB58 prophage genes during inter-species bacterial interactions.

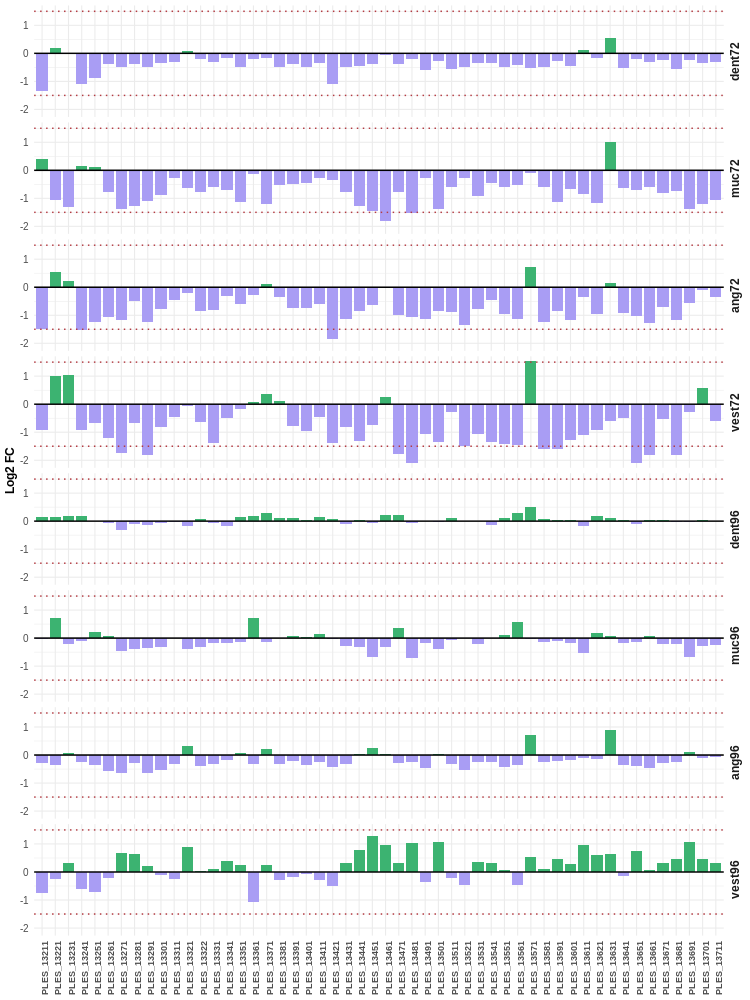

**Supplementary Figure 3.** Prophage 3: F10-like. Log2FC values distribution of LESB58 prophage genes during inter-species bacterial interactions.

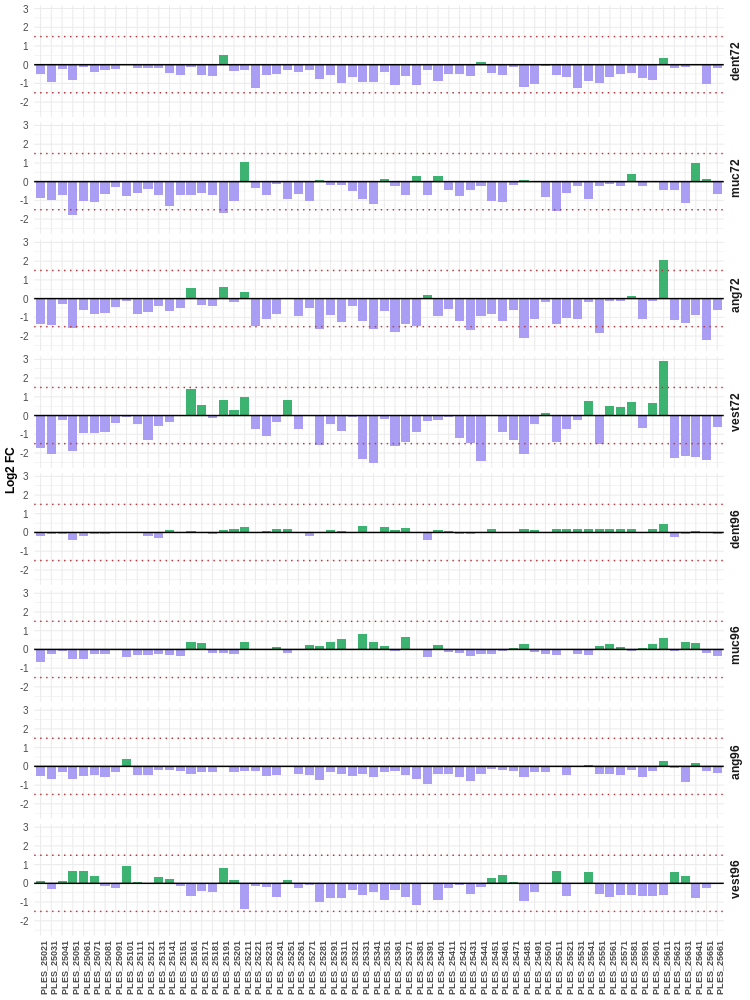

**Supplementary Figure 4.** Prophage 5: D3-like. Log2FC values distribution of LESB58 prophage genes during inter-species bacterial interactions
